# Selective spread of mobile antibiotic resistance genes in wastewater microbiomes driven by the non-antibiotic pharmaceutical carbamazepine

**DOI:** 10.1101/2025.09.09.675050

**Authors:** Eda Deniz Erdem, Bing Li, Thomas U. Berendonk, Uli Klümper

## Abstract

Carbamazepine (CBZ), a widely used anticonvulsant, is a persistent aquatic micropollutant that withstands biodegradation and accumulates in wastewater-impacted environments. While non-antibiotic pharmaceuticals, including CBZ, have been shown to stimulate horizontal gene transfer (HGT) under laboratory conditions, their effects in complex microbial communities remain poorly understood. Here, we applied an experimental evolution framework to wastewater microbiomes exposed to a gradient of environmentally relevant CBZ concentrations (0–50 µg/mL) for three days. CBZ reduced growth of the community in a concentration-dependent manner. Still, community composition was largely unaffected by CBZ exposure. However, antibiotic resistance gene (ARG)- and mobile genetic element (MGE)-specific responses emerged. Among the tested 26 ARGs, four clinically relevant ones (*bla*_CMY_, *bla*_OXA-48_, *bla*_CTX-M_, *dfr*A1) increased in relative abundance in a dose-dependent manner, correlating with proliferation of IncP and IncW plasmids and the transposon IS26, pointing to enhanced HGT as the underlying mechanism. In contrast, only *erm*F abundance rose independently of MGEs, suggesting direct selection. Other ARGs and MGEs, including integron integrase *int*I1 and IncQ plasmids, showed no consistent CBZ response. Effects were strongest at subinhibitory concentrations (0.05-5 µg/mL), matching upper levels detected in surface waters. These findings demonstrate that CBZ can selectively promote AMR dissemination through both HGT and direct selection, but in a targeted rather than community-wide manner. Our results highlight the need to consider non-antibiotic pharmaceuticals in AMR surveillance and environmental risk assessment frameworks, as they may shape the environmental resistome beyond antibiotic exposure.

## 1. Introduction

Antimicrobial resistance (AMR) among human and animal pathogens poses a mounting global health threat (WHO, 2015). In 2021, bacterial AMR was linked to approximately 4.7 million deaths worldwide, including 1.1 million directly caused by resistant infections, with projections estimating that by 2050, AMR could be directly responsible for 1.9 million deaths annually, and 8.2 million deaths associated with it [1,2]. A sizeable share of mitigation efforts regarding the proliferation of antibiotic-resistant bacteria (ARB) and antibiotic resistance genes (ARGs) have been focused on clinical and veterinary settings [3]. Despite this, there is a growing consensus on the necessity of understanding how the environment at large acts as a reservoir and conduit for AMR [3–6].

One of the key mechanisms underlying the rapid spread of AMR is horizontal gene transfer (HGT), enabling bacteria to acquire resistance genes from other strains or species [7–9]. Conjugation, the transfer of plasmids among bacteria, plays a major role, as high proportions of ARGs are plasmid-borne [10,11]. Conjugative plasmid transfer in natural microbial communities typically occurs at low frequencies due to fitness costs associated with plasmid carriage and the low likelihood of encounters between donor and permissive recipient bacteria under environmentally realistic cell densities [12,13]. Despite these generally low transfer rates, a variety of biotic and abiotic factors can significantly alter conjugation frequencies in environmental microbiomes in either direction, thereby shaping the dissemination of AMR [14]. Such factors include heavy metals [9], temperature shifts [15], and various xenobiotic compounds such as microplastics [16,17], nanoparticles [18,19], and complex mixtures such as cigarette smoke [20]. Microbial interactions themselves, such as competition or cooperation, can also modulate plasmid transfer frequencies positively or negatively [21–23]. So can exposure to antibiotics, which in addition serves as a selective force on those plasmids encoding ARGs [24].

More recently, attention has turned to non-antibiotic pharmaceuticals, which are increasingly recognized as important environmental drivers of HGT. These compounds are widespread in aquatic ecosystems due to continual discharge from households, hospitals, agriculture, and pharmaceutical production [25,26]. While microbial responses to antibiotics have been studied extensively, the impact of non-antibiotic pharmaceuticals remains a newer and rapidly evolving field, with recent focus on effects on selection for AMR [27] and especially horizontal gene transfer [14,28]. Emerging evidence shows that even pharmaceuticals lacking antimicrobial activity can enhance plasmid transfer by interacting with bacterial physiology. However, most studies to date have relied on simplified laboratory systems, often using single donor strains and high plasmid loads under conditions that diverge substantially from natural environments.

The anticonvulsant carbamazepine (CBZ), widely used in human medicine to treat epilepsy and mood disorders, represents a valuable model compound for investigating how non-antibiotic pharmaceuticals influence microbial communities and the environmental resistome. Due to its tricyclic structure, CBZ persists against biodegradation and is poorly removed during conventional wastewater treatment processes [29]. Coupled with its frequent and consistent consumption, this persistence leads to accumulation in aquatic environments, with concentrations reaching up to 3600 ng/L in surface and groundwaters [30].

A conjugation-promoting effect of CBZ was first discovered in 2019 using single-strain mating assays between *Escherichia coli* and *Pseudomonas putida* [31]. The underlying mechanisms were identified as enhanced reactive oxygen species (ROS) production, increased membrane permeability, and subsequent higher expression of plasmid-borne transfer-related genes. Although further experiments extended these findings to complex microbial recipient communities, plasmid transfer was still assessed from an artificially introduced single donor strain carrying the RP4 plasmid at unnaturally high abundances, thus limiting ecological realism [32].

These findings raise an important question: Do the plasmid transfer effects of non-antibiotic pharmaceuticals observed under laboratory conditions translate to environmental contexts? Especially, the ability of CBZ to promote the spread of natural ARG-encoding plasmids has yet to be tested in environmental bacterial communities, where such plasmids are present at far lower abundances. Moreover, whether the detected plasmid transfer promotion shown for IncP-1-type plasmids [31,32] is consistent across the diverse types of environmental plasmids remains unknown. Finally, if beyond stimulating horizontal gene transfer, CBZ, similar to other non-antibiotic pharmaceuticals [27], may directly or co-select for specific ARGs, has not been explored. Such questions can best be answered using experimental evolution approaches, where microbial communities are under controlled conditions exposed to gradients of the compound in question.

Two recent studies, one using fish microbiota [33] and one using anaerobic digestion of activated sludge [34], observed first indications of concentration- and duration-specific increases in ARG levels under CBZ exposure. Both studies focused on a limited number of ARGs and tested limited or difficult-to-translate concentration ranges of CBZ while also not targeting plasmids, and were thus not able to offer further insights into the research questions posed above.

To overcome these limitations and assess the environmental relevance of CBZ exposure on AMR dynamics, we applied an experimental evolution framework using a complex wastewater microbial community. Communities were evolved for three days under a gradient of six CBZ concentrations. We then quantified changes in the absolute and relative abundance of 26 ARGs and five MGEs, including markers for three of the most common plasmid incompatibility groups found in environmental microbiomes. Using this approach, we show that CBZ exposure is associated with gene- and plasmid-specific shifts in ARG and MGE relative abundance, highlighting the need to carefully consider effects of non-antibiotic pharmaceuticals in environmental AMR risk assessments.

## 2. Materials & Methods

### 2.1. Wastewater effluent community

The complex microbial community originated from the treated effluent of the Kreischa wastewater treatment plant (Dresden, Saxony, Germany) that was collected in June 2023. Two litres of the sampled effluent were filtered through 0.22 μm pore-size sterilized filters (Millipore). Three out of six filters used were placed in 50 mL tetrasodium pyrophosphate (TSPP) buffer and vortexed to facilitate the detachment of microorganisms. The remaining filters were set aside for DNA extraction. The TSPP buffer was then added to 150 mL of LB broth in its entirety, along with the filters, to avoid a major loss of microbiome. This mixture was incubated for 24 hours at 30°C to obtain a liquid overnight culture of the microbial community to be used for evolution experiments and DNA extraction. The resulting DNA samples were denoted as Day 0 samples.

### 2.2. Evolution experiments under carbamazepine presence

Evolution experiments were conducted in 12-well cell culture plates over three days with daily passages. On Day 0, the 12-well plates were prepared with LB growth medium containing six concentrations of CBZ (Sigma-Aldrich, St. Louis, MO, USA) in a gradient (0, 0.05, 0.5, 5, 50, and 500 µg/mL), and the overnight complex microbial culture was added in a 1:100 dilution after washing with PBS and re-suspension in fresh LB medium. CBZ concentrations were selected to incorporate environmentally relevant concentrations along with those included in previous studies [30,31]. The range of the gradient was further expanded to 500 µg/mL to investigate the possible antibacterial activity of CBZ and the dose-dependent response of the microbial community. Six biological replicates were conducted for each concentration and incubated at 30°C.

Since the CBZ stock solution was prepared at 50 mg/mL using DMSO, the replicate cultures prepared for 50 µg/mL and 500 µg/mL concentrations had final DMSO concentrations of 1% and 10%. DMSO can cause inhibition of bacterial growth at these concentrations [35]. Therefore, solvent controls were established where four replicates of both DMSO concentrations were represented to differentiate the true effect of CBZ and the potential impact of DMSO at such high concentrations with regard to the structural change of the microbiome.

Every 24 hours, the bacterial cultures were transferred to a new well containing fresh LB medium with the corresponding CBZ or DMSO concentration in a 1:100 dilution to replenish nutrients. After 72 hours, samples were collected from all replicates, and the DNA extraction protocol was initiated immediately. Samples were named based on the CBZ (CBZ_0_, CBZ_0.05_, CBZ_0.5_, CBZ_5_, CBZ_50_, CBZ_500_) or DMSO (DMSO_1%_, DMSO_10%_) concentrations the communities were evolved at and are subsequently referred to using this nomenclature.

### 2.3. DNA extraction

All DNA extractions were carried out using the DNeasy PowerSoil Pro Kit (Qiagen, Hilden, Germany) per the manufacturer’s protocol. In the cases where DNA was extracted from bacterial cultures, 2 mL of liquid culture at carrying capacity was collected, followed by centrifugation, a wash step, and the final resuspension in PBS. The isolated DNA was evaluated for its quantity and quality using a NanoDrop™ (ThermoScientific, Waltham, MA, USA) and stored at -20°C for further analyses.

### 2.4. 16S rRNA qPCR

To assess if overall growth patterns of the community were affected by CBZ presence during the three-day experimental period, 16S rRNA gene copy numbers (gene copies/ng of DNA) in the samples were determined with SYBR Green-based quantitative PCR and used as an indicator of total bacterial abundance. A final concentration of 300 nM of the primer pair 338F (CCTACGGGAGGCAGCAG) - 518R (ATTACCGCGGCTGCTGG) [36] was used along with 10 µL of Luna® Universal qPCR Master Mix (New England Biolabs, Massachusetts, USA) and 5 µL of template DNA extract at 2 ng/µL in a final volume of 20 µL per reaction. A recombinant plasmid [37] was extracted and used as the qPCR standard at target concentrations of 10^8^ - 10^2^ copies per reaction. For each DNA sample, three technical replicates were set up in Real-Time qPCR in a C1000 Touch™ Thermal Cycler (BioRad, Hercules, CA, USA), and the cycling conditions were 44 cycles of 15 s at 95°C and 30 s at 60°C, preceded by an initial denaturation cycle of 95°C for 10 min. DNA isolates from treated wastewater effluent and initial augmented culture were analysed as well to assess the effect the pre-experiment protocol had on the bacterial load. Only the results without multiple peaks in the melting curve analysis and where the standard curve amplification efficiency was 0.9–1.1 and R² ≥ 0.99 were accepted. Ten-fold dilution of template DNA revealed no PCR inhibition.

The obtained 16S RNA gene copy numbers were used to calculate the number of cell divisions that took place within the communities during the evolution experiment, which was X in the following equation:

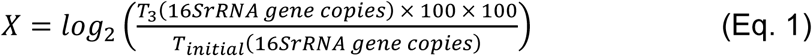

### 2.5. Identification of Microbial Community

The composition and taxonomic diversity of pre-evolved and evolved microbial communities were evaluated by Illumina MiSeq amplicon sequencing of the 16S rRNA gene. DNA samples at 10 ng/µL were transferred to the IKMB sequencing facility (Kiel 458 University, Germany), where the bacterial 16S rRNA gene was sequenced using primers targeting the V3-V4 region (V3F: 5′-460 CCTACGGGAGGCAGCAG-3′ V4R: 5′-GGACTACHVGGGTWTCTAAT-3). Molecular analysis of the resulting sequences was carried out using the Mothur software package v.1.48.0 [38] per MiSeq SOP [39]. Subsequently, OTUs were generated at 97% sequence similarity and taxonomically classified based on the RDP classifier [40]. Sequencing depth of the samples varied between 403338 reads at the highest and 4896 at the lowest; this minimum value was then used for subsampling. Differences in the microbial composition of the samples were first analysed on the phylum level. Thereafter, OTU-based beta diversity of the sequences was assessed via non-metric multidimensional scaling (NMDS) by the Bray-Curtis dissimilarity index. The analysis of molecular variance (AMOVA) function of Mothur was used to determine the statistical significance of the spatial clustering of the samples, as indicated.

### 2.6. High-throughput quantitative PCR

Samples were also used to quantify the possible spread of ARGs and MGEs in the microbial communities evolved under CBZ pressure and the pre-experimental microbial culture. DNA was sent to Resistomap Oy (Helsinki, Finland) to determine the relative abundances of a selection of 26 ARGs and 5 MGEs, along with the 16S rRNA gene, which was used for normalization (Table 1). ARGs were selected to cover different antibiotic classes and genes of clinical and environmental relevance. Among the 5 MGEs, special emphasis was placed on the three markers corresponding to three different incompatibility groups of broad host range plasmids, IncP, IncQ, and IncW [41–43], as CBZ-based induction of plasmid transfer was previously observed for at least IncP-1 plasmids from single donor strains [31,32]. Further, transposable element IS*26* and class 1 integron integrase gene *intI1* were considered to be proxy targets for remaining plasmid families due to their ubiquitous nature and close association with ARG-carrying plasmids [44–46].

**Table 1.**
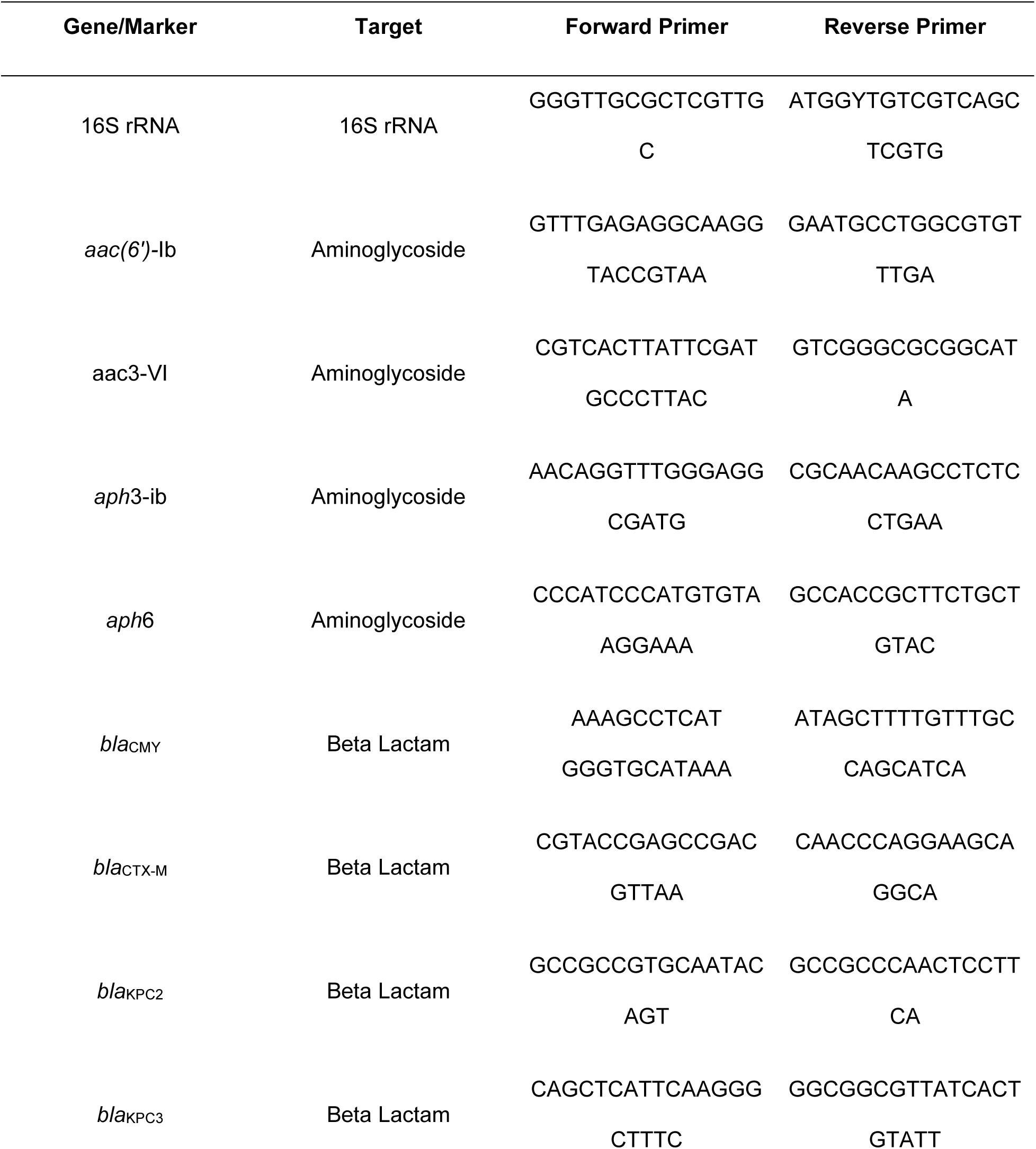
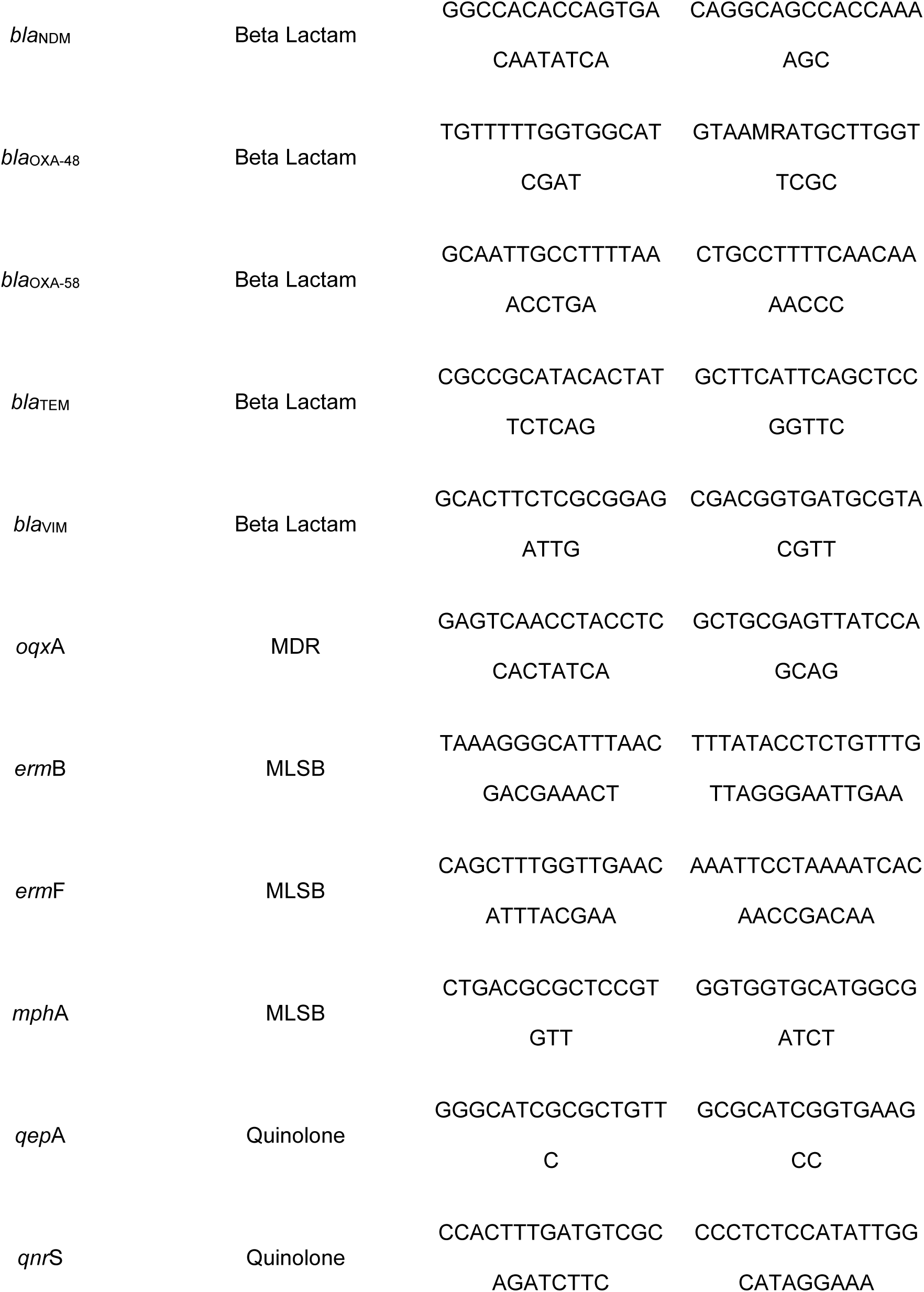
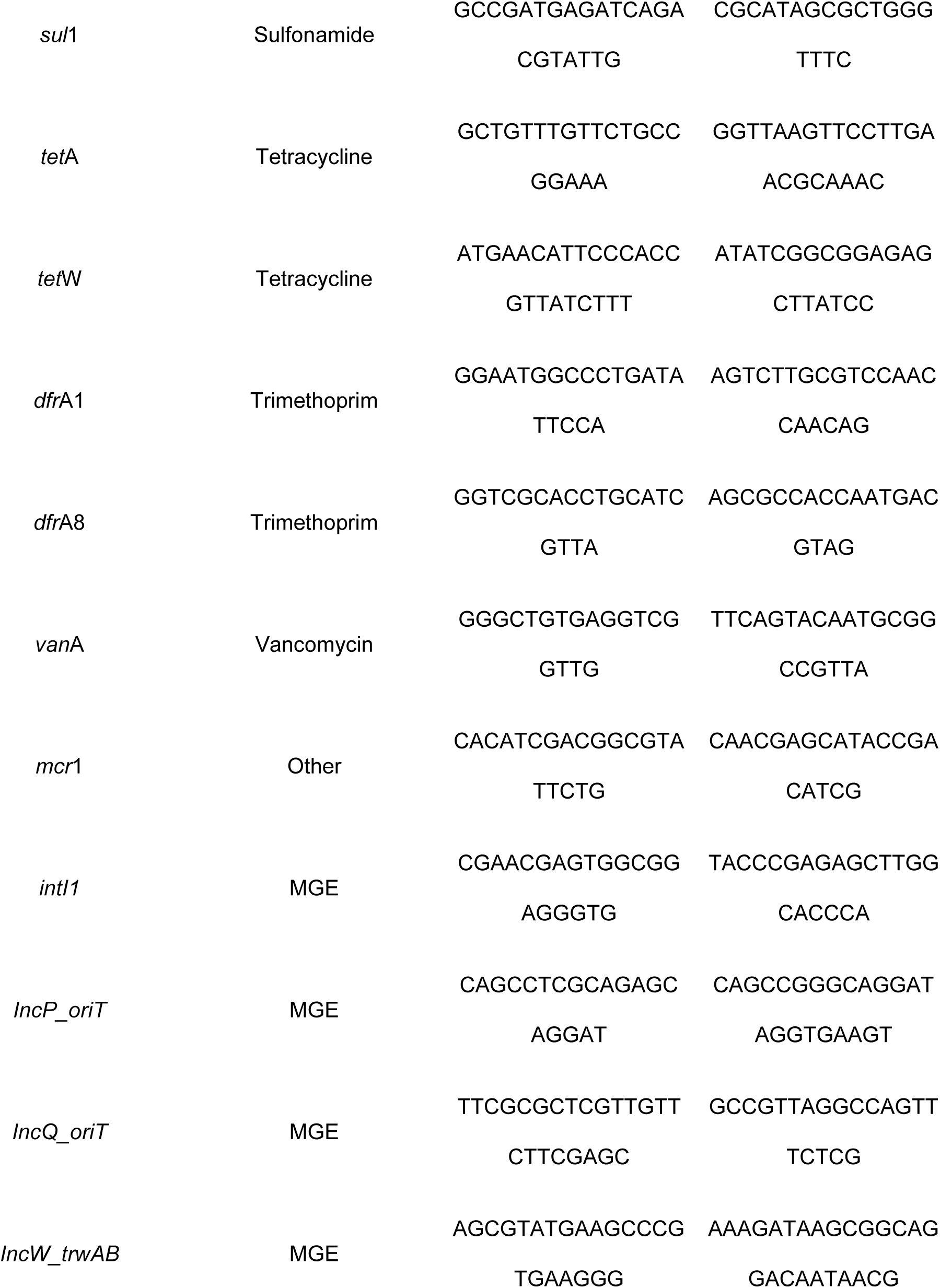
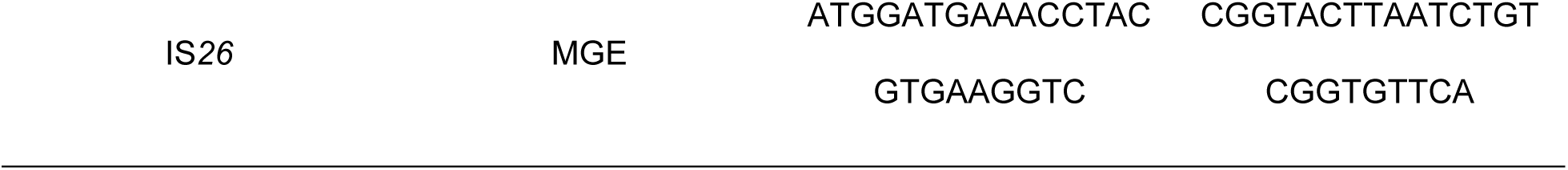
qPCR primers used in this study based on Stedtfeld et al. [76].

The HT-qPCR protocol included the preparation of 100 nL of PCR reaction mixture with SmartChip TB Green Gene Expression Master Mix (Takara Bio, Japan), nuclease-free PCR-grade water, 300 nM of each primer, and 2 ng/μL DNA template. Cycling conditions were an initial denaturation step at 95°C for 10 min, after which 40 cycles of 95°C for 30 s and 60°C for 30 s followed. For each primer set, a melting curve analysis was performed. For the analysis of the resulting data, cycle thresholds (CT) above 31 were considered below the detection limit, and replicates where efficiencies were not between 1.8 and 2.2 were filtered out. The relative copy number of genes of interest normalized to the 16S rRNA gene was calculated using the ΔCT method [47], where CT refers to the mean CTs of three technical replicates.

### 2.7. Statistical analysis and data visualization

All statistical tests, data manipulations, and visualizations were executed in R Studio v2024.04 [48]. Prior to statistical analyses, the normalcy of data was checked by the Shapiro-Wilk test along with equality of variances, and the appropriate statistical test to use was determined based on the results.

All correlation analyses were performed using the Spearman rank correlation method. When required, a log_10_ transformation of CBZ concentration was used, with CBZ_0_ set as one log_10_-fold change below CBZ_0.05_. Table 2 provides a classification of the correlation strength we used in this study according to [49]. All pairwise comparisons were done with the Mann-Whitney U test, and the Kruskal-Wallis rank sum test was performed for multiple comparisons. Unless stated otherwise, P-value adjustments were performed using the Benjamini and Hochberg false discovery rate method provided by the multtest R package [50], and the threshold for the P-value for significance was 0.05. Dose-response curves and correlograms were produced with the drc and corrplot R packages, respectively [51,52]. All remaining plots were created within the ggplot2 package [53].

**Table 2.** Strength of Spearman’s rank correlation as described by Kuckartz et al. [49].

| Magnitude of $R_s$ | Strength of correlation |
| --- | --- |
| $R_s > 0.70$ | Very strong correlation |
| $0.50 < R_s < 0.70$ | Strong correlation |
| $0.30 < R_s < 0.50$ | Moderate correlation |

The rate of increase coefficient (RoI) of a target gene at CBZ concentration *i* was calculated as described previously [54], and used to infer the increase in the relative abundance of a gene from the beginning of the evolution experiment, normalized by the shift that occurred without CBZ influence:

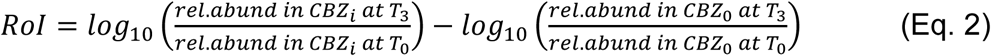

## 3. Results

### 3.1. Minor impact of CBZ exposure on microbial community composition

To examine the capacity of CBZ to facilitate the spread of AMR by altering horizontal gene transfer and selection dynamics in a complex bacterial community, an experimental evolution experiment with minimal manipulations was performed. A wastewater microbial community was exposed to six concentrations of CBZ (0, 0.05, 0.5, 5, 50, 500 µg/mL) or their relevant solvent controls for three days with daily passages to replenish nutrients and CBZ after the communities reached carrying capacity. First, the impact of CBZ on evolved community composition was elucidated via 16S rRNA gene-based amplicon sequencing.

All communities underwent significant phylogenetic changes during the evolution experiment compared to the pre-evolved initial community (Figure 2A, F = 18.52, p < 0.0001, AMOVA with Bonferroni correction for multiple testing). Three distinct clusters of the evolved communities were observed: The first cluster consisted of the communities treated with CBZ at the highest concentration of 500 µg/mL (Figure 2B, initial vs. CBZ_500_: F = 20.60, p < 0.05), while the second cluster entailed the communities evolved in the absence of CBZ (control) as well as all those evolved at the remaining CBZ concentrations (0.05 – 50 µg/mL). Each treatment in this cluster was grouped significantly away from the initial samples (Figure 2B, F = 14.55 - 25.44, all p < 0.05) and the CBZ_500_ endpoint communities (Figure 2B, F = 27.19 - 49.62, all p < 0.05). The final cluster consisted of the solvent control corresponding to the CBZ_500_ treatment (DMSO_10%_), which caused a significant dissimilarity to the aforementioned cluster of CBZ_0-50_ (F = 30.58, p < 0.001). While this cluster also displayed significant divergence from the CBZ_500_ cluster, the effect size was low (F = 8.52, p = 0.03), indicating high similarity in the diversity of the evolved samples and that the increased solvent concentration could have mainly contributed to any potentially observed effects in the CBZ_500_ samples. Contrarily, as the DMSO_1%_ grouped with the CBZ_0-50_ cluster (F = 4.41, p > 0.05), no significant solvent effects at that concentration were observed. Thus, the CBZ_500_ and DMSO_10%_ samples were excluded from further analyses, and only the CBZ_0-50_ cluster was analysed further. Within this primary cluster, disparities between communities evolved at the different CBZ concentrations were observed. Namely, CBZ_0_, CBZ_0.05_ and CBZ_0.5_ samples were significantly separated from CBZ_5_ and CBZ_50_, with no significant differences within these groups (Figure 2B, CBZ_0_ vs. CBZ_5_: F = 5.12; CBZ_0_ vs. CBZ_50_: F = 10.84; CBZ_0.05_ vs. CBZ_50_: F = 7.06; CBZ_0.5_ vs. CBZ_5_: F = 3.04; CBZ_0.5_ vs. CBZ_50_: F = 6.89, all p < 0.05).

**Figure 1:**
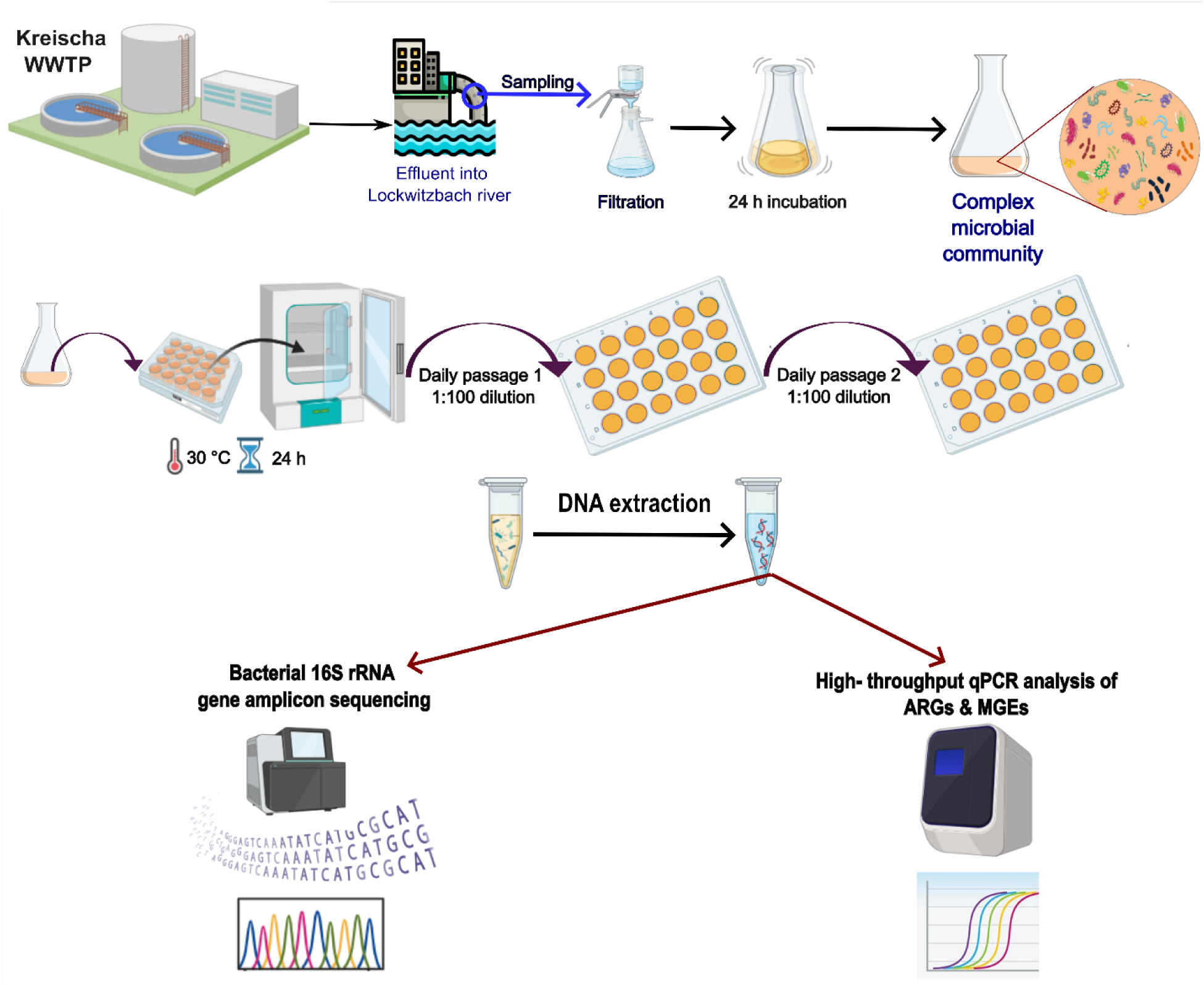
Experimental evolution framework and methods used in this study. The effluent of the WWTP was sampled to generate the model complex microbial community, which was then evolved with varying CBZ concentrations. Resulting DNA isolates were used to identify changes to the microbial community compositions and ARG and MGE relative abundances in endpoint communities.

**Figure 2:**
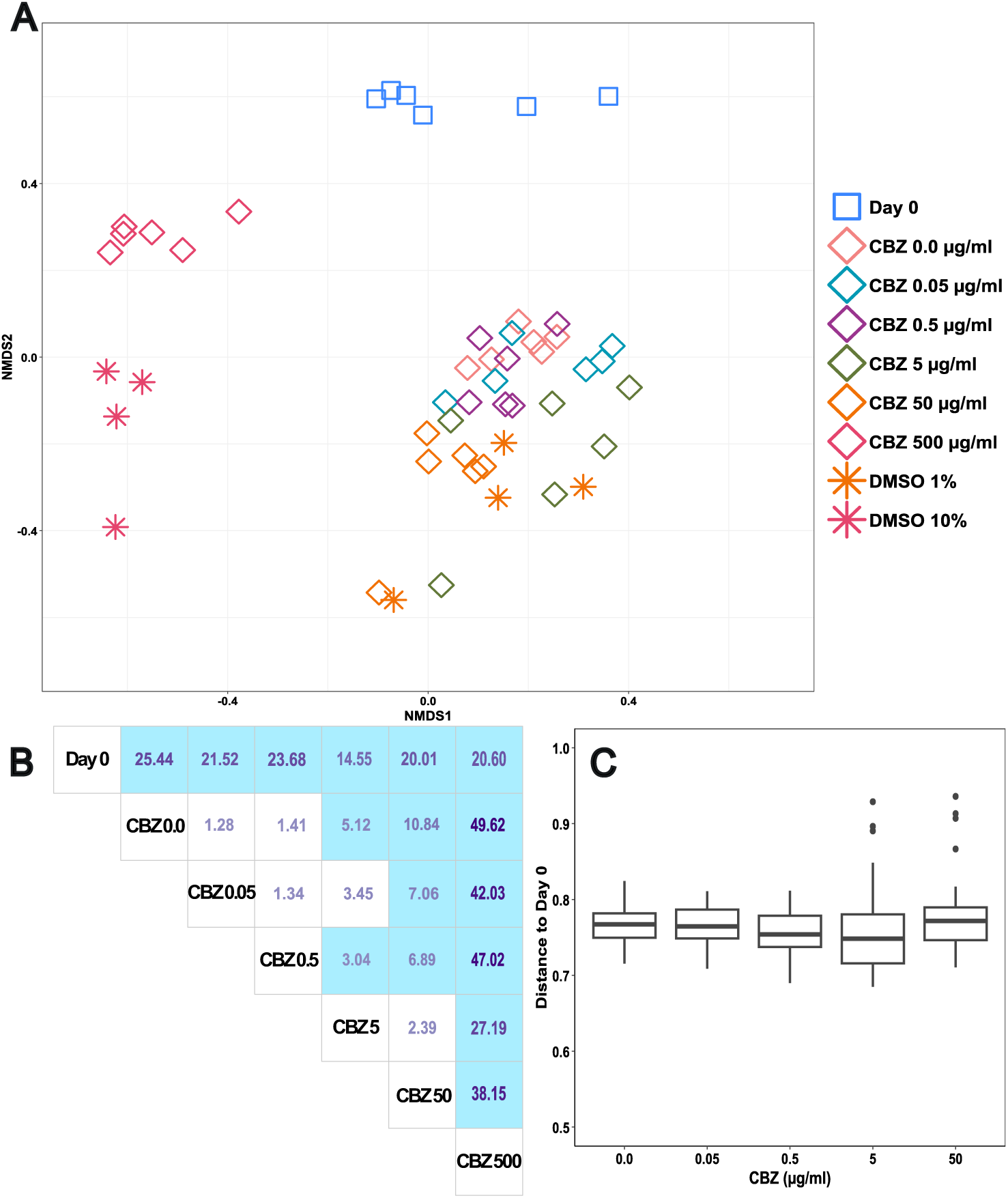
Beta-diversity analysis of the microbial communities. OTU-based beta diversity microbial communities before (Day 0) and after (CBZ_0-500_, DMSO_1%_ and DMSO_10%_) the evolution experiment as indicated by the Bray-Curtis dissimilarity index. (A) NMDS plot visualising the phylogenetic variations between microbial communities. (B) AMOVA post-hoc pairwise comparisons of the initial microbial community and samples evolved under CBZ presence, given as F-statistics. Significance is indicated as blue background (p < 0.05). (C) Bray-Curtis dissimilarity of CBZ_0-50_-evolved microbial communities to the initial community.

Nevertheless, the effect sizes of these variations in community compositions between endpoint samples were much smaller than those with the pre-experimental culture of Day 0. Moreover, the CBZ concentration did not correlate with Bray-Curtis dissimilarity between endpoint communities and those of Day 0 (Figure 2C, R_S_ = 0.06, p = 0.42, Spearman rank correlation).

### 3.2. CBZ did not affect community diversity, but slightly reduced growth

11461 OTUs were identified across all samples. At the phylum level, Proteobacteria (56.38 ± 23.21%), Firmicutes (35.56 ± 17.32%), and Bacteroidetes (6.72 ± 6.29%) were the most abundant phyla in the initial microbial communities (Figure 3A). After evolution, Fusobacteria and Bacteroidetes significantly rose in prominence by 24 (p = 0.0001, Mann-Whitney U test) and 1.8 (p = 0.03) times compared to the initial levels (Figure 3A). Contrarily, two of the major phyla, Proteobacteria and Firmicutes, did not undergo significant changes in average relative abundance. Similarly, no significant differences among the CBZ_0-50_ endpoint samples were detectable on the phylum level (Figure 3A). While the evolution of the communities had clear effects on the beta-diversity as well as the phylum-level distribution, only minor effects of the CBZ concentration on the diversity of the communities were noticeable.

**Figure 3:**
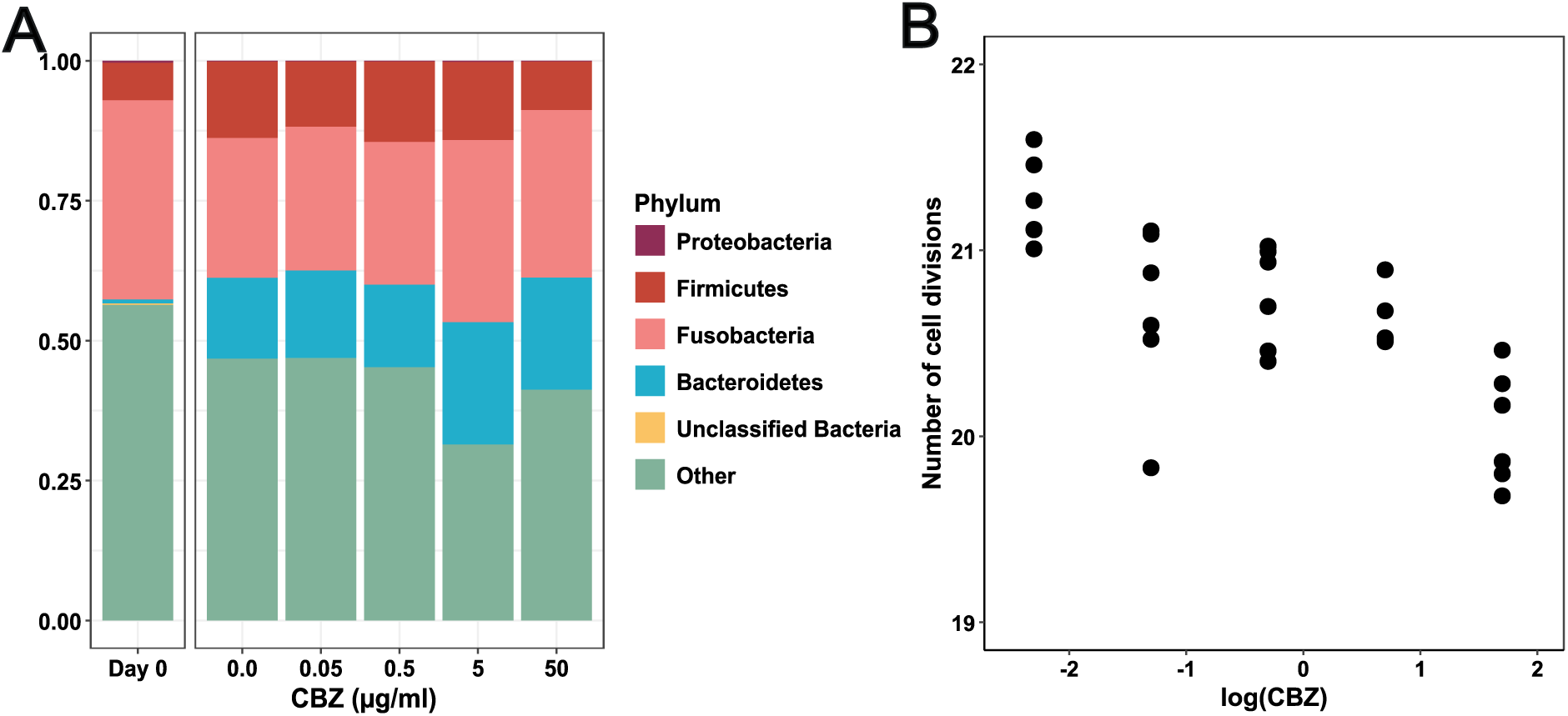
Effect of CBZ presence on the phylum-level composition and bacterial growth in the microbial communities. (A) Proportions of dominant phyla in the microbial communities. “Others” refers to phyla whose average relative abundance is less than 1%. (B) Number of cell divisions (Eq. 1) in the evolved communities plotted against CBZ concentration (log_10_ basis).

However, CBZ exposure reduced growth during the evolution experiment. Cell divisions undergone by the CBZ_0-50_ endpoint communities showed a significant negative correlation with CBZ concentration (R_S_ = -0.78, p < 0.001, Spearman rank correlation, Figure 3B). However, this reduction in growth was only minor, with a difference of approximately a single cell division less at the highest CBZ concentration compared to the non-exposed control. This indicates that while CBZ did not affect overall community composition, it slightly impaired growth dynamics. Still, with this only minor growth effect and as community composition remained stable across all evolved communities, relative rather than absolute abundances of ARGs and MGEs were subsequently utilised.

### 3.3. Increasing CBZ concentrations are linked to higher total relative plasmid abundance

Based on the minor effects CBZ exposure had on evolved community composition on growth, whether it altered the relative abundance of 26 ARGs and 5 MGEs through either selection or modulated horizontal gene transfer rates was tested. The total normalised relative abundance of tested ARGs decreased significantly by 52.27% during the evolution experiment in the absence of CBZ (Figure 4A, initial vs. CBZ_0_: p = 0.01, Mann-Whitney U test). Contrarily, the total relative plasmid marker abundance (0.002 ± 0.0001) was not significantly impacted by the evolution experiment in the absence of CBZ (Figure 4B, p = 0.38). For the remaining MGEs, transposable element IS*26* and class 1 integron integrase gene *intI1,* an even sharper reduction of 88.43% and 35.43% was observed (Figure 4C & D, both p < 0.01).

**Figure 4:**
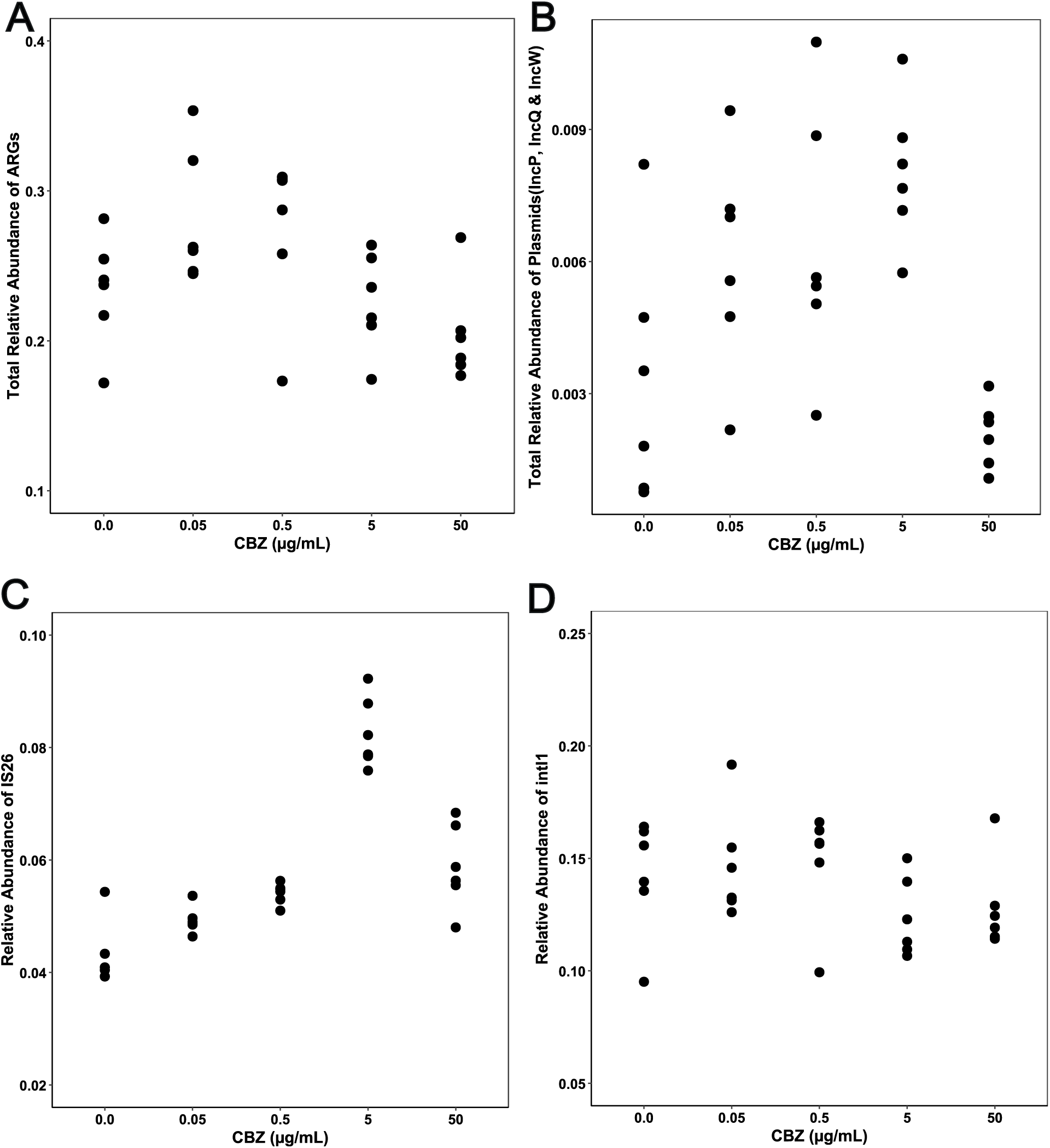
Relative abundances of the genes of interest by type. Total relative abundances of 26 ARGs (A), IncP, IncW, and IncQ plasmid markers (B), transposable element IS*26* (C), and class 1 integron integrase gene *intI1* (D) in the evolved microbial communities as a function of increasing CBZ concentrations.

Among the evolved communities, no significant changes in relative abundance of total ARGs, total plasmids, or other MGEs were observed for any individual CBZ concentration compared to the non-exposed control (CBZ_0_ vs. CBZ_0.05-50_: all p>0.05, Mann-Whitney U test, FDR-adjusted). Moreover, no dose-response correlation of CBZ for either of these abundances was detected when considering all CBZ concentrations except for IS*26* (Total ARGs: R_S_ = -0.30, p = 0.11; total plasmids: R_S_ = 0.01, p = 0.97; *intI1*: R_S_ = 0.21, p = 0.26; IS*26*: R_S_ = 0.75, p < 0.0001; Spearman rank correlation). However, at the CBZ concentration range between 0 and 5 µg/mL, a strong and significant dose-dependent relationship was observed between CBZ exposure and total relative abundance of plasmid markers exclusively (R_S_ = 0.58, p = 0.003). Despite plasmids as the main vehicles of ARG spread correlating with CBZ in this concentration range, no correlation with total ARGs (R_S_ = -0.05, p = 0.84) and no correlation between total relative plasmid and ARG abundance in the dataset was detected (R_S_ = 0.25, p = 0.19). This could be, as only a minor proportion of the tested ARGs are actually plasmid- or generally MGE-bound and hence potentially affected by CBZ-induced higher transfer rates, or that only individual ARGs are selected for by CBZ. Consequently, ARG, plasmid, and IS*26* dynamics across the CBZ gradient on a gene-by-gene basis were investigated next. As Class 1 integrons (intI1) were not affected by CBZ at any concentration, these were excluded from subsequent analysis.

### 3.4. Gene-dependent dose-response effects of CBZ exposure

To determine whether CBZ-induced spread of the 26 individual ARGs (conferring resistance to 10 different classes of antibiotics) occurred and if it was connected to the induction of plasmid or IS transfer, their fluctuations in relative abundances were evaluated subsequently. The number of detected ARGs was comparable in all evolved microbial communities (24.31 ± 0.70 ARGs, p = 0.57, Kruskal-Wallis rank sum test), with only one ARG (*bla*_OXA-58_) discarded from in-depth analysis due to being under the detection limit for the majority of samples.

Evolution alone, independent of CBZ pressure, had a sizable impact on the ARG and MGE relative abundance. 16 out of 25 ARGs, 2 out of the 3 plasmid markers, and IS*26* all decreased significantly in abundance after evolution compared to the initial sample (FDR-adjusted p = 0.003 - 0.001, Mann-Whitney U test, Figure 5A). Only one ARG, *erm*F, which confers resistance against macrolides, had a 2.3-fold elevated significant abundance after evolution (FDR-adjusted p = 0.003).

**Figure 5:**
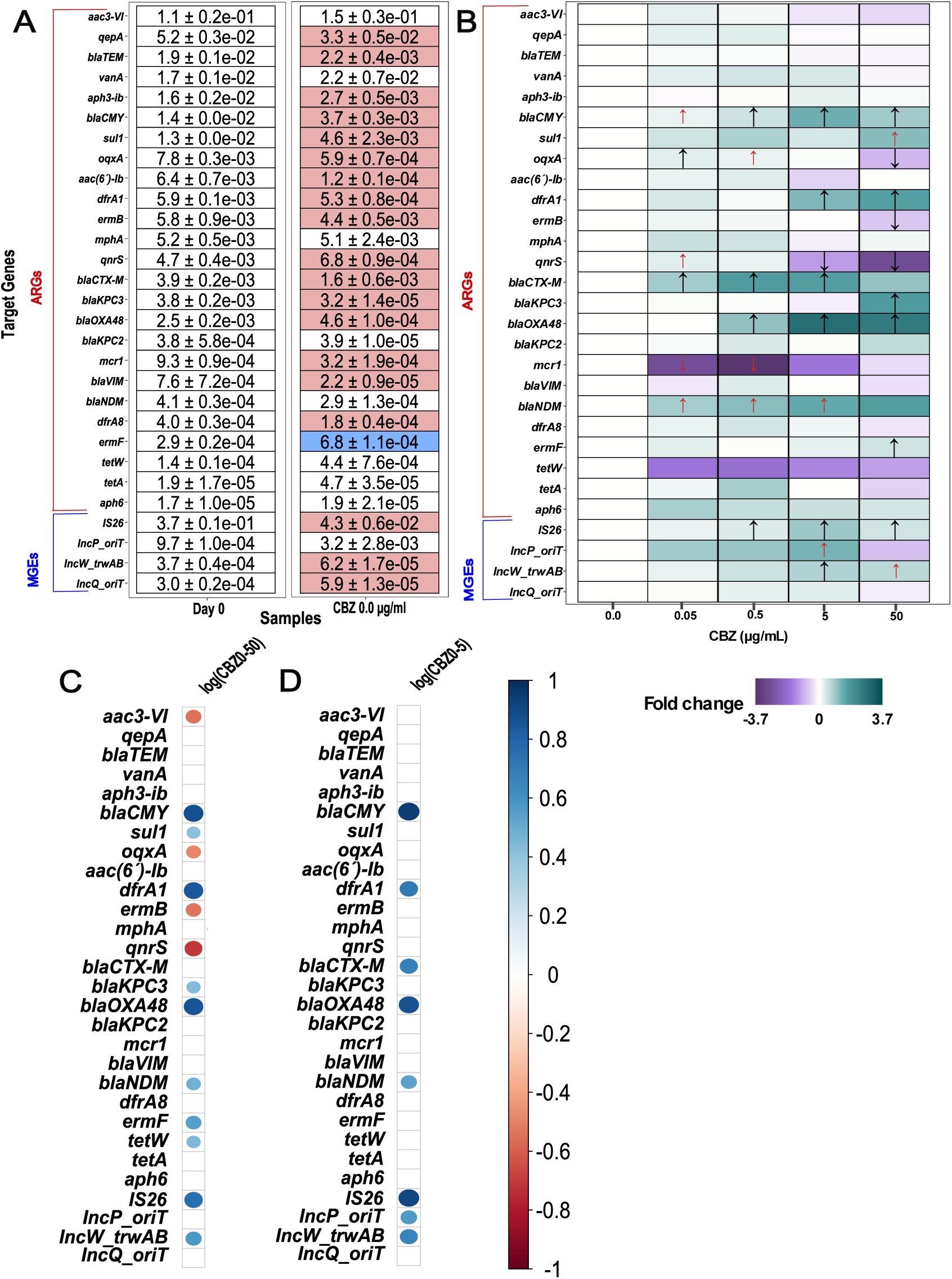
Impact of CBZ exposure on relative abundances of individual genes of interest. (A) Changes in relative abundances of ARGs and MGEs during evolution without CBZ presence. Purple background on CBZ_0_ samples refers to a significant decrease from the relative abundance of the gene in initial community (Day 0) while a significant increase is referred to as blue (FDR-adjusted p < 0.05, Mann-Whitney U test). (B) Heatmap of relative abundances of ARGs and MGEs of interest in the evolved communities, based on log2 fold change from CBZ_0_ samples. Arrows indicate a significant shift in relative abundances of the genes compared to that in the CBZ_0_ communities (FDR-adjusted p < 0.05 for black arrows, 0.05 < FDR-adjusted p < 0.10 for red arrows, Mann-Whitney U test). Spearman correlation analyses between relative abundances of target genes and log10 basis of full (C) and intermediate (D) range of CBZ concentration. Only significant correlations are shown (FDR-adjusted p < 0.05). For the entire figure, target genes are ordered based on their relative abundances in the initial microbial community from highest to lowest.

In the evolved communities, CBZ had varying effects on ARG and MGE relative abundances (Figure 5B). For 13 of 25 ARGs, no significant difference compared to the control was detected at any concentration (all FDR-adjusted p > 0.1, Mann-Whitney U test). These included the 5 most abundant ARGs: *aac3*-VI, *qep*A, *bla*_TEM_, *vanA,* and *aph*3-ib. Only 5 of the 25 ARGs exhibited significant relative abundance surges across the majority of CBZ-exposed communities (log_2_-fold change: 0.21-2.84, FDR-adjusted p = 0.02-0.06, Mann-Whitney U test). Among these were four β-lactam ARGs (*bla*_CTX-M_, *bla*_CMY_, *bla*_OXA-48_, *bla*_NDM_) and *dfr*A1 that confers resistance to trimethoprim. Additionally, significant increases at only the highest CBZ concentration compared to the control were detected for three ARGs (*bla*_KPC3_, *erm*F, *sul*1). Among the remaining 4 ARGs, significant decreases in abundance were detected only once at low (*mcr*1) or once at the highest CBZ concentration (*erm*B) compared to the control. Initial increases at low and sharp decreases at high CBZ concentrations were detected twice (*qpx*A, *qnr*S, Figure 5B).

Across the tested MGEs, significant increases in relative abundance across the majority of CBZ-exposed communities were observed for IncW plasmids and IS*26* (log2-fold change: 0.33-0.94, FDR-adjusted p = 0.02-0.06, Mann-Whitney U test). For IncP plasmids, an increase at intermediate CBZ concentrations (FDR-adjusted p = 0.07) followed by a 1.6-fold, but non-significant decrease in relative abundance at the highest concentration was detected, mirroring the previous trend for total relative plasmid abundance. Unlike for IncW and IncP plasmids, no effects on IncQ plasmids at any concentration were observed (FDR-adjusted p = 0.33-0.90).

In the absence of a comprehensive trend for the spread of ARGs and MGEs in evolved microbial communities, Spearman’s rank correlation analysis helped clarify the role of CBZ concentration-dependent dose-responses. When taking all CBZ concentrations from 0 to 50 µg/mL into account, significant positive correlations with the relative abundances of 8 of the 25 ARGs were identified (Figure 5C). Except for *bla*_CTX-M_, these included all ARGs for which previous increases in relative abundance were detected at several concentrations, as well as all ARGs for which significant increases were only detected at the highest CBZ concentration and *tet*W, for which no individual sample displayed any significant change in abundance (Figure 5C). The strongest positive dose-response relationships were found for *bla*_CMY_, *bla*_OXA-48_, and *dfr*A1 (R_S_ = 0.84 - 0.87, FDR-adjusted p < 0.05), while the remaining positive correlations with CBZ concentration for *erm*F, *sul*1, *bla*_KPC3_, *bla*_NDM_, and *tet*W were moderate (R_S_ = 0.42 - 0.47, FDR-adjusted p < 0.05). Negative correlations were identified between CBZ concentrations and the relative abundances of the four genes *aac3*-VI, *oqx*A, *erm*B, and *qnr*S (R_S_ = 0.49-0.71, 0.0001 < FDR-adjusted p < 0.02). Among the MGEs, very strong (R_S_ > 0.70) and moderate (0.30 < R_S_ < 0.50) positive correlations with CBZ concentration were detected for IS*26* and IncW plasmids, respectively.

A second Spearman rank correlation analysis was performed within the range of CBZ concentrations that had previously yielded a significant relationship with total relative plasmid abundance (CBZ 0–5 µg/mL, Figure 5D). This removed the majority of moderate correlations, except for *bla*_NDM_ (R_S_ = 0.53, FDR-adjusted p = 0.03), including all negative correlations of ARGs. All previous strong correlations remained significant (*bla*_CMY_, *bla*_OXA-48_, and *dfr*A1, IS*26*, and IncW plasmids, R_S_ = 0.67-0.95, FDR-adjusted p < 0.005), whereas two new strong correlations between CBZ and *bla*_CTX-M_ as well as IncP plasmids were apparent (R_S_ = 0.68 & 0.57, FDR-adjusted p < 0.05).

As concentration-dependent increases for a number of ARGs and MGEs were observed, next their dependency on each other was tested.

### 3.5. Correlations with MGEs explain CBZ dose-response effects on ARGs

To investigate whether the observed CBZ dose-response increases in certain ARGs are based on HGT, correlations between MGE and ARG relative abundance within the evolved microbial communities using the full (0-50 µg/mL, Figure 6A) and the intermediate (0-5 µg/mL, Figure 6B) CBZ concentration range were tested.

**Figure 6:**
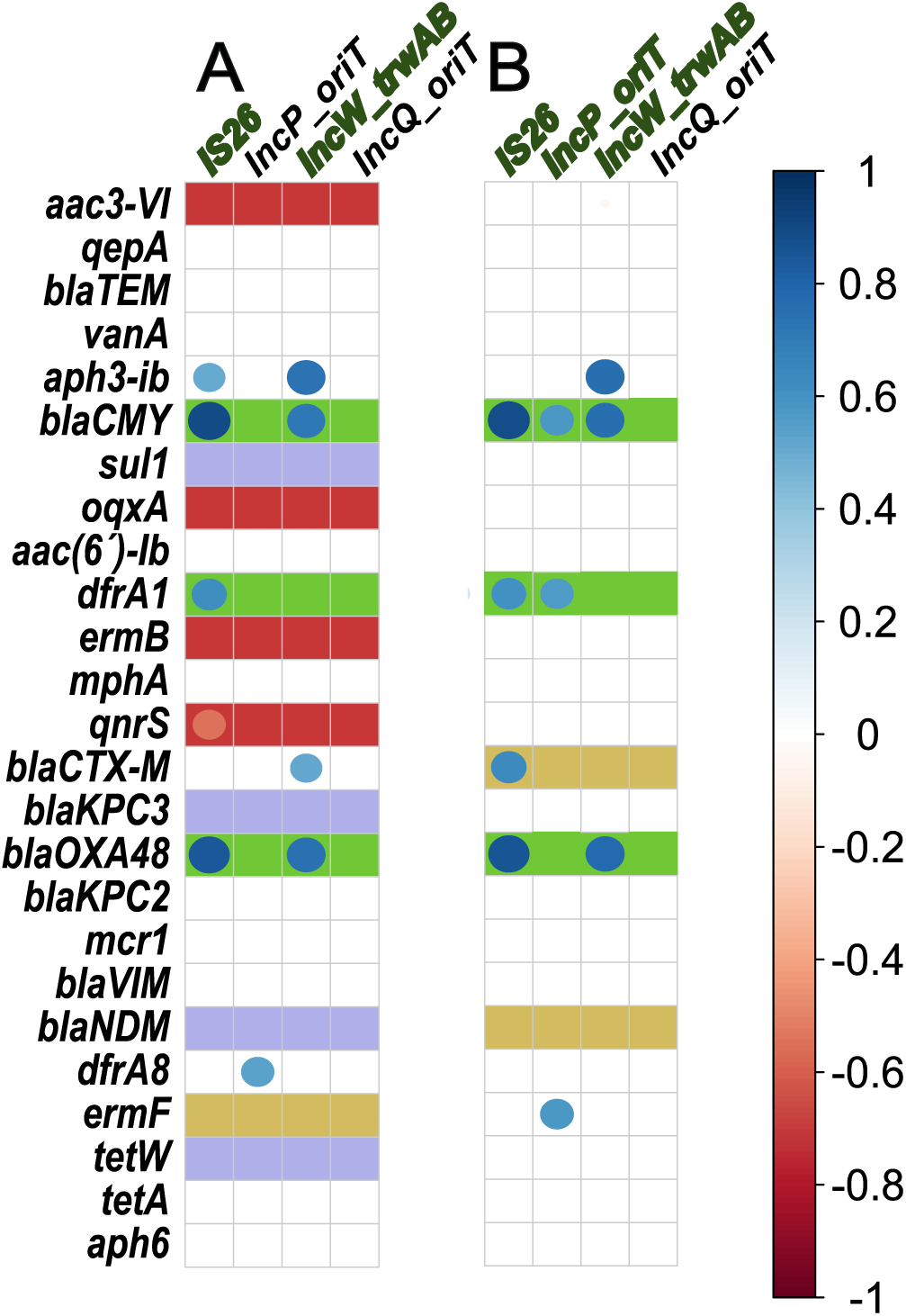
Correlations between the relative abundances of ARGs and MGEs. Spearman’s correlation analyses were performed with relative abundances from the entire (A, 0-50 µg/ml) and intermediary (B, 0-5 µg/ml) CBZ concentration span. Only significant correlations are shown on correlograms (FDR-adjusted p < 0.05). Strong correlations between CBZ concentration and relative abundances of MGEs (described in Figure 5 C & D and categorised by [49] are indicated as green legend text. Strengths of correlation between the relative abundance of ARGs and CBZ concentration (described in Figure 5C&D) are given as background colour codes of green (very strong correlations, R_S_ > 0.7), yellow (strong correlations, 0.5 < R_S_ < 0.7), purple (moderate correlations, 0.35 < R_S_ < 0.5) and red (negative correlations) as categorised by [49].

In the intermediate CBZ concentration dataset, 4 out of 5 ARGs with strong and very strong positive correlations to CBZ concentrations (Table 2, R_S_ > 0.68, green and first yellow background in Figure 6B) correlated with at least two of the 3 MGEs (IncP plasmids, IncW plasmids, IS*26*) that also displayed positive correlation with CBZ. *bla*_CMY_ positively correlated with all 3 MGEs (R_S_ = 0.58-0.89, FDR-adjusted p = 0.00001-0.04, *drf*A1 correlated with IncP plasmids and IS*26* (R_S_ = 0.56 & 0.61, FDR-adjusted p < 0.05) and *bla*_CTX-M_ and *bla*_OXA-48_ that correlated with IncW plasmids and IS*26* (R_S_ =0.55-0.86, FDR-adjusted p = 0.0001-0.048). Contrarily, *bla*_NDM_ that displayed a less strong correlation with CBZ (Second yellow background in Figure 6B) did not correlate with any of the MGEs. Among the 20 ARGs that did not correlate with CBZ concentrations, in general, no positive correlations with the relative abundance of any MGEs were detected. Exceptions were *aph*3-ib, which encodes an aminoglycoside-modifying enzyme and correlated with IncW plasmids and IS*26* (R_S_ = 0.75 & 0.55, respectively, FDR-adjusted p < 0.05), and *erm*F, which displayed a relatively weak correlation with IncP plasmid abundance (R_S_ = 0.57, p = 0.04). IncQ plasmids that were not affected by CBZ concentrations did not display a single ARG correlation throughout.

Similar trends were visible when investigating the full dataset, with those ARGs displaying very strong positive correlations with CBZ (green background) generally displaying MGE correlations, while those with only strong positive (yellow), moderate positive (purple), no (colourless), or negative correlations with CBZ mainly displayed no MGE correlations (Figure 6A). However, in the full dataset, again, a few correlations between MGEs and ARGs that had no CBZ correlations (*aph*3-ib, *van*A, *dfr*A8 & *tet*A) were detected. These could originate from dose-response curves reaching a plateau at lower concentrations, meaning that investigating a higher concentration range adds additional randomness in the correlation analysis. Consequently, their rate of increase during the evolution experiment (Eq. 2) was modelled as a function of CBZ dose-response (Figure 7).

**Figure 7:**
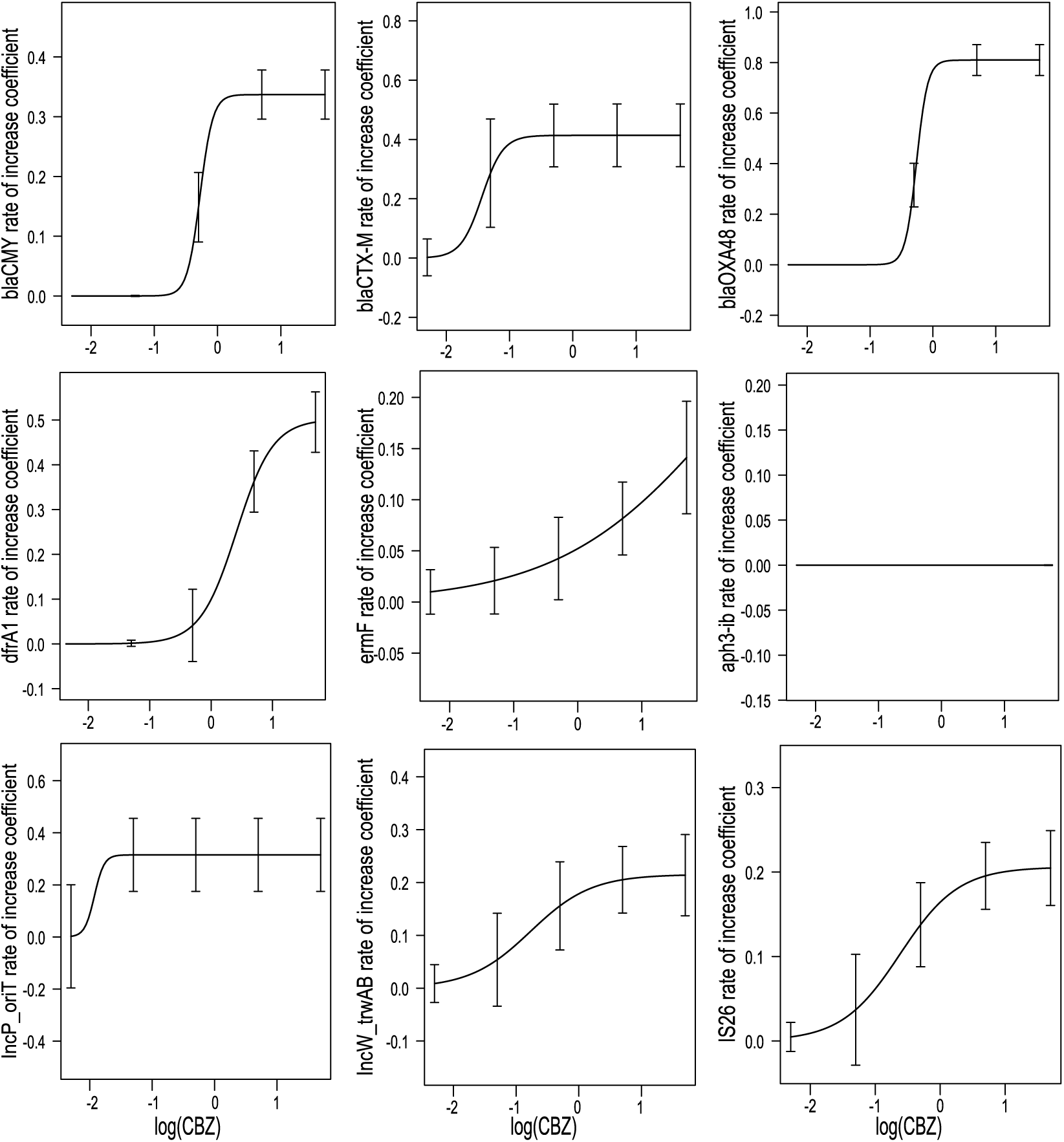
Dose-response dynamics between rate of increase and CBZ concentration for selected genes. Dose-response models built to examine the rate of increase of relative abundances (Eq. 2) of *bla*_CMY_, *bla*_OXA-48_, *bla*_CTX-M_, *dfr*A1, *erm*F, *aph*3-ib, IncP, IncW, and IS*26* markers in relation to the log10 basis of CBZ concentration. The three-parameter log-logistic (LL.3) function from the drc R package [51] was used to build the models and the error bars represent model-based standard errors.

Indeed, for IncP plasmids, a plateau of the rate of increase was reached at a very low dosage of CBZ (between 0.05 and 0.5 µg/mL). Contrarily, no plateau for the rate of increase was apparent for IS*26* and IncW plasmids, although the model implicates that the maximum response may be observed around the maximum tested CBZ concentration of 50 µg/mL. For the 4 ARGs that displayed strong CBZ dependence and strong MGE correlations throughout (*bla*_CMY_, *bla*_OXA-48_, *bla*_CTX-M_ & *drfA*1), clear dose-response relationships were modelled. For *bla*_CTX-M_, a similarly low plateau concentration of CBZ to that observed for IncP plasmids (between 0.05 and 0.5 µg/mL) was observed. For the *bla*_CMY_ and *bla*_OXA-48_ genes, the plateau concentration was in the intermediate range between 0.5 and 5 mg/mL, while for *dfr*A1, similar to IS*26* and IncW plasmids, no plateau for the rate of increase was apparent with the model implicating that the maximum response may be observed around the maximum tested CBZ concentration of 50 µg/mL. For those ARGs with potentially random correlations with MGEs but not with CBZ (Figure 6B), dose-response modelling was either impossible (*aph*3-ib) or relative abundances were increasing throughout with a steep slope, even as a response to the highest CBZ concentration tested (*erm*F) (Figure 7).

## 4. Discussion

Previous laboratory studies have shown that CBZ can promote plasmid-mediated ARG transfer, with single-strain and community-level mating assays reporting up to 12- and 7.1-fold increases in conjugation frequency, respectively, for the model plasmid RP4 [31,32]. In parallel, two recent studies using more complex community settings [33,34] observed first indications of concentration- and duration-specific increases in ARG levels under CBZ exposure, suggesting potential selection or co-selection effects. Our experimental evolution study builds on this foundation by testing these mechanisms under more environmentally realistic conditions. We found that, despite a minor, dose-dependent growth-inhibiting effect, CBZ had no major effect on community structure or overall relative ARG abundance. Rather, it selectively increased the relative abundance of specific ARGs and MGEs at subinhibitory and environmentally relevant concentrations (0–5 µg/mL). Four clinically relevant ARGs, *bla*_CMY_, *bla*_OXA-48_, *bla*_CTX-M_, and *dfr*A1, displayed clear dose-dependent increases correlating with the proliferation of IncP and IncW plasmids and the transposable element IS*26*, pointing to enhanced horizontal gene transfer as the likely mechanism. However, other tested MGEs were not significantly affected by CBZ exposure. ARGs whose abundance did not correlate with any MGE abundance were, throughout, not significantly affected by CBZ. The lone exception was *erm*F, which increased independently of MGE dynamics, indicating a role for direct selection. Together, our results suggest that CBZ can influence AMR propagation in complex microbial communities through both HGT and gene-specific selection, albeit in a more nuanced and limited manner than laboratory studies might imply.

Significant correlations were observed between CBZ concentration and the relative abundances of transposable element IS26 and the conjugative plasmids IncP and IncW. Whereas previous studies reported strong CBZ-induced effects on IncP plasmids such as RP4 [31,32], we observed that effects on IS*26* abundance were even greater than on IncP plasmid abundance. This is likely due to IS*26*’s frequent association with conjugative plasmids [46], which benefit from enhanced transfer under CBZ exposure [28,31,32]. Moreover, once present in a host, IS*26* can amplify within cells through replicative transposition, especially under stress conditions, independent of horizontal transfer [44,46]. This dual capacity to spread both between and within cells may explain the particularly strong IS*26* response to CBZ.

These observations are consistent with the proposed mechanism of CBZ-induced ROS overproduction, which activates the bacterial SOS response, thereby promoting expression of conjugation-related genes such as *tra* via derepression of *kor*AB regulators in IncP and potentially IncW plasmids [28,55–60]. Additionally, IS*26* activity may also be directly enhanced by the same ROS-triggered SOS stress cascade, as transposase expression and replicative transposition are known to respond to such stress [44]. The strongest correlations between CBZ exposure and MGE and MGE-associated ARG relative abundance were observed within the environmentally relevant 0-5 µg/mL range. This concentration range (0–5 µg/mL) reflects the upper levels of CBZ detected in aquatic environments, including effluents, surface water, and occasionally groundwater [26,29,30]. Associations weakened across the full 0-50 µg/mL gradient used in this study, suggesting a potential plateau or even inhibitory effect at higher CBZ concentrations. Similar non-linear dynamics have been reported in earlier studies [28,31,61]. ROS production and conjugation or transformation efficiencies increased at low CBZ concentrations but failed to rise further or even declined at higher doses. Moreover, in the case of transformation efficiencies, subinhibitory concentrations of non-antibiotic pharmaceuticals most effectively enhanced HGT, while excessive doses yielded diminished effects [61]. Together, these findings support a model in which moderate ROS stress promotes plasmid transfer and MGE activation, but excessive stress may suppress these processes via cellular damage, defensive regulatory systems, or metabolic burden. However, these promoting effects appear to be MGE-specific: while IncP, IncW, and IS*26* responded clearly, mobilizable IncQ plasmids, despite their broad host range, showed no significant dose-dependent change. This is unlikely to result from a lack of compatible conjugative plasmids, needed to support IncQ transfer [62], as, for example, IncP and IncW plasmids were shown as abundant, and due to the natural origin of the community, likely the same is true for other conjugative plasmid groups. Instead, it suggests that CBZ-induced ROS preferentially stimulates self-transmissible conjugative systems, whereas mobilisation-dependent elements such as IncQ [63,64] are less affected in complex communities.

Similarly, Class 1 integron integrase gene *intI1* did not increase in response to CBZ, despite being under *lex*A repression and part of the SOS regulon. This lack of response may reflect a regulatory activation threshold for *intI1* that was not met under CBZ-induced ROS stress. While moderate ROS levels may have been sufficient to activate *tra* operons and IS*26* transposition, previous studies show that *intI1* requires strong, sustained SOS induction, typically triggered by DNA-damaging antibiotics like fluoroquinolones [65–67]. Moreover, *intI1* is not self-transmissible, relying on its association with conjugative elements for horizontal transfer, and in natural communities, a substantial proportion of class 1 integrons are chromosomally encoded [68]. This chromosomal integration may decouple integron dynamics from plasmid-level conjugation effects, masking any impact of CBZ on *intI1* abundance.

Overall, these findings emphasise that the impact of CBZ on ARG propagation is gene- and MGE-specific, rather than broad or uniform. While ROS-mediated stimulation of conjugation and transposition appears to explain much of the observed HGT activity, selective pressures should not be ruled out. Notably, the resistance gene *erm*F exhibited a clear dose-response increase, yet did so independently of known MGEs, suggesting that direct selection by CBZ or adaptation to lab conditions may have driven its proliferation. Similar selection dynamics for resistance determinants have been observed with other non-antibiotic compounds [27]. Distinguishing between CBZ’s effects on conjugation versus selection remains a key challenge for future studies.

This study builds on previous work by extending the investigation of CBZ-driven antimicrobial resistance dynamics from simplified laboratory setups to a complex, natural microbial community, while applying environmentally relevant concentrations of CBZ across a broad gradient. Whereas earlier studies mainly focused on single-strain conjugation assays [31], relied on artificially introduced plasmids and donors [32], or focused on limited targets and concentrations [33,34], our experimental evolution approach provides a more ecologically realistic model to assess ARG and MGE behaviour in response to CBZ exposure in situ. By quantifying both plasmid and transposon dynamics, our study advances understanding of how non-antibiotic pharmaceuticals influence horizontal gene transfer of ARGs in natural microbial communities. Nonetheless, several limitations should be acknowledged. First, our experiment was conducted over a short duration and under nutrient-rich laboratory conditions that do not fully reflect the complexity and resource limitations of natural ecosystems. Subtle effects of low, environmentally relevant CBZ concentrations may only become apparent over longer periods. Indeed, a recent aquatic mesocosm study demonstrated that community-level changes may take weeks to months to manifest at µg/L CBZ levels [69]. Extended exposures under more representative environmental conditions are hence needed to better capture the long-term ecological and evolutionary impacts of CBZ on MGE and ARG proliferation. Thus, the limited effects observed at the lower concentrations used in this study likely represent a conservative baseline, potentially underestimating the cumulative risk posed by persistent pharmaceutical pollution.

Second, we inferred resistance dissemination from shifts in the relative abundance of ARGs and MGEs and their correlations, without resolving specific gene-host relationships. This limits our ability to distinguish between mechanisms such as horizontal transfer, clonal expansion, or intracellular amplification. To improve resolution, future research should incorporate techniques such as long-read metagenomics, Hi-C sequencing [70], epicPCR [71] or multiplex ddPCR analysis [72] that allow host- and MGE-associations of ARGs.

Despite these limitations, our findings raise broader questions about the evolutionary pressures that non-antibiotic pharmaceuticals may impose on natural microbial communities. For example, if long-term CBZ exposure could drive co-selection and co-evolution between ARGs and MGEs such as IncP, IncW, or IS*26*, by selectively enriching those ARG vehicles best suited to thrive under chemical stress, remains to be investigated. If so, this could shape the structure of the environmental resistome in ways not visible in short-term assays. Our results also suggest that, in addition to conjugative plasmids, MGEs like IS*26*, due to their strong and mechanistically grounded response to CBZ exposure, may serve as biomarkers of pharmaceutical-driven HGT and be useful for AMR surveillance in wastewater-impacted ecosystems.

## 5. Conclusion

Our study demonstrates that the non-antibiotic pharmaceutical carbamazepin can selectively promote the spread of antimicrobial resistance genes ARGs in complex microbiomes. Whereas overall community composition remained largely unaffected, CBZ exposure drove dose-dependent increases in specific clinically relevant ARGs such as *bla*_CMY_, *bla*_OXA-48_, *bla*_CTX-M_, and *dfr*A1. These increases were linked to IncP/IncW plasmids and the transposon *IS*26, pointing to enhanced horizontal gene transfer as a central mechanism. In contrast, the macrolide resistance gene *erm*F rose independently of MGEs, indicating a role for direct ARG selection.

These findings highlight that CBZ, and potentially other non-antibiotic pharmaceuticals [28,31,32,34], can shape the environmental resistome in a targeted rather than community-wide manner, with concentrations in the range of those detected in surface waters exerting the strongest effects.

Given the widespread occurrence of CBZ and other non-antibiotic pharmaceuticals in aquatic environments [26] and the here and elsewhere [28,31,32,34] demonstrated capacity to modulate resistance gene dynamics, we thus recommend their inclusion in AMR risk assessment frameworks and environmental monitoring programs [4,73,74]. However, unlike for antibiotics, where defining predicted no-effect concentrations for selection (PNEC_res_) is straightforward [75], defining such PNECs for compounds that mainly affect AMR through HGT modulation or co-selection remains more challenging. Still, surveillance systems should be expanded to consider not only antibiotic usage but also the broader pharmaceutical context that shapes resistance dissemination in real-world microbial ecosystems to allow mitigation efforts to be established.

## Acknowledgements

The authors thank Steffen Kunze for logistical support during sampling.

## 8. Data Availability

The datasets supporting the conclusions of this article are included within the article and its additional files or are available through the corresponding author upon reasonable request. The original sequencing data are available in the NCBI sequencing read archive under project accession number PRJNA1321686.

## 9. Author Contributions

**EDE:** Conceptualisation, Methodology, Validation, Formal analysis, Investigation, Data Curation, Writing - Original Draft, Writing - Review & Editing, Visualization; **BL:** Writing - Review & Editing, Project administration, Funding acquisition; **TUB:** Conceptualisation, Resources, Writing - Review & Editing, Supervision, Project administration, Funding acquisition; **UK**: Conceptualisation, Methodology, Resources, Data Curation, Writing - Review & Editing, Visualization, Supervision, Project administration, Funding acquisition

